# Segmental outflow and trabecular meshwork stiffness in an ocular hypertensive mouse model

**DOI:** 10.64898/2026.02.03.703547

**Authors:** Cydney A. Wong, A. Thomas Read, Guorong Li, Amia Loveless, Nina Sara Fraticelli-Guzmán, Andrew J. Feola, Todd Sulchek, W. Daniel Stamer, C. Ross Ethier

## Abstract

**Purpose:** Elevated intraocular pressure (IOP) due to increased outflow resistance through the trabecular meshwork (TM) is a major risk factor for primary open-angle glaucoma. Outflow through the TM is segmental, consisting of high flow (HF) and low flow (LF) regions. Here, we investigate how ocular hypertension impacts segmental outflow using a dexamethasone (DEX) mouse model and compare TM stiffness between HF and LF regions.

**Methods:** Nanoparticles containing DEX or vehicle were injected twice weekly in 2–4-month-old C57BL/6J mice (n=14), and IOP was measured weekly. At week 4, mouse eyes were perfused *in vivo* with fluorescent nanospheres to assess flow patterns and the circumferential percentage of high, intermediate, and low flow regions in each eye. Sagittal sections were collected from HF and LF regions, and atomic force microscopy (AFM) was used to measure tissue stiffness. Immunofluorescent labeling was used to compare fibronectin and α-SMA protein levels.

**Results:** DEX treatment significantly elevated IOP by an average of 33.3% and altered tracer distribution but not the percentage of HF and LF regions around the circumference. No significant differences in TM stiffness were detected between DEX-treated and control mice, or between HF and LF regions. Increased fibronectin in LF regions of DEX-treated eyes suggested subtle TM structural changes that were not detected by AFM.

**Conclusions:** Dexamethasone alters segmental flow distribution and may impact cell contractility rather than ECM stiffness to cause IOP elevation in young mice. These findings better characterize the nature of segmental outflow and TM mechanics in this model of steroid-induced glaucoma.

## 1. Introduction

Ocular hypertension, or elevated intraocular pressure (IOP), is the only modifiable risk factor for Primary Open-Angle Glaucoma (POAG)^1^. In healthy eyes, IOP is maintained by homeostatic processes involving the production and drainage of aqueous humor from the anterior chamber of the eye. Aqueous humor drains primarily through the conventional outflow pathway, which consists of the trabecular meshwork (TM), Schlemm’s canal (SC), and collector channels/aqueous veins. In patients with ocular hypertension, there is increased resistance to outflow through the conventional pathway, with the juxtacanalicular (JCT) region of the TM as well as the SC inner wall being the main sources of resistance to aqueous humor outflow^2–5^.

The biomechanical properties of the trabecular meshwork (TM) and the inner wall of Schlemm’s canal (SC) play an important role in influencing aqueous humor outflow and IOP. In glaucomatous human eyes and in animal models of ocular hypertension, the TM becomes stiffer^6–8^, and this increased TM stiffness is correlated with increased outflow resistance^9^. Further, pharmacological approaches that reduce TM stiffness, such as rho-kinase inhibitors effectively lower IOP^10,11^.

Importantly, aqueous humor outflow is not uniformly distributed around the circumference of the eye. In both healthy and ocular hypertensive eyes, outflow through the TM is segmental, i.e., there are distinct high flow (HF) and low flow (LF) regions. This non-uniformity in outflow has been demonstrated using a variety of tracer perfusion studies^12–18^ where a greater concentration of tracer indicates a higher flow. Previous work has shown that LF regions have lower outflow facility (i.e. higher outflow resistance)^19^, and that viable human TM cells extracted from HF and LF regions show differences in stiffness; specifically, cells from LF regions of glaucomatous eyes are particularly stiff compared to cells from HF regions and compared to cells from normotensive eyes^20^. Differences in various extracellular matrix (ECM) components and ECM regulators between HF and LF regions have also been identified via transcriptomic and proteomic analysis in human TM^18,21–23^. Consistent with these findings, *ex vivo* human TM tissue is stiffer in LF regions than in HF regions, and this disparity is amplified following exposure to elevated perfusion pressure, which induces further stiffening in LF regions^23^. Additionally, ECM produced by LF-derived TM cells is intrinsically stiffer than that produced by HF-derived cells, and treatment with dexamethasone further enhances ECM stiffness in both regions^24^. Collectively, these studies indicate that regional differences in TM biomechanics are strongly associated with segmental outflow patterns; however, these differences have mainly been characterized in human tissue. In mice, which are widely used as animal models of ocular hypertension and POAG, differences between biomechanical properties of HF and LF regions have not been well-characterized.

Glucocorticoids, such as dexamethasone (DEX), are commonly employed to induce ocular hypertension in animal models^25^, yet it is not known whether DEX differentially alters the biomechanics of HF and LF regions. Establishing whether segmental flow patterns and segmental differences in TM stiffness are altered in mice under ocular hypertensive conditions is therefore critical for understanding the translational relevance of this model. Thus, the objective of this study is to characterize segmental flow distribution in a dexamethasone-induced mouse model of ocular hypertension and use atomic force microscopy (AFM) force mapping to assess TM stiffness in HF and LF regions of control and dexamethasone-treated mouse eyes, providing new insight into the biomechanical basis of segmental outflow regulation in this ocular hypertension mouse model.

## 2. Methods

### 2.1. Experimental Design Overview

We used an established mouse model in which DEX is delivered via periocular injection to induce ocular hypertension in mice^26,27^. After IOP elevation was observed, we perfused a fluorescent tracer into the eyes in vivo to identify high and low flow regions. The circumferential tracer distribution pattern in the limbus was imaged, and then sagittal cryosections from both HF and LF regions of the eye were collected for both stiffness measurements by AFM and immunofluorescent staining.

### 2.2. Nanoparticle synthesis

Nanoparticles containing dexamethasone (DEX-NPs) and control nanoparticles (CON-NPs) were prepared in Dr. Stamer’s laboratory at Duke University as previously described^28^. Briefly, penta block (PB) co-polymer (PDLLA-PCL-PEG-PCL-PDLLA, AK099) was purchased from Akina, PolySciTech (West Lafayette, IN, USA). To prepare DEX-NPs, the PB polymer was mixed with dexamethasone acetate (PHR1572, Sigma Aldrich) at a 20:1 ratio by weight. CON-NPs were prepared in the same way without adding dexamethasone. Both DEX- and CON-NPs were then dissolved in ethyl acetate and eluted dropwise in a 0.1% vitamin E solution. After sonication, the solution was then transferred to a 0.3% vitamin E solution. The solvent was evaporated in the fume hood overnight at room temperature stirring at 400 rpm.

After ultracentrifugation and two additional washes with ddH₂O for further concentration and purification, DEX-NPs or CON-NPs were resuspended in PBS to a final NP concentration of 1mg/20μL, vortexed for 10 min, and then sonicated for 10 min before injection.

### 2.3. Animals, IOP Measurement, and Tracer Perfusion

All animal procedures were approved by the Institutional Animal Care and Use Committee (IACUC) of Duke University and Georgia Institute of Technology and were consistent with the ARVO Statement for the Use of Animals in Ophthalmic and Vision Research. C57BL/6J wild-type mice were purchased from the Jackson Laboratory (Bar Harbor, Maine, USA), bred and housed in clear cages, and kept in housing rooms at 21°C on a 12:12 h light:dark cycle. 2–4-month-old males (n=14) received bilateral 20 μL subconjunctival or periocular injections of nanoparticles loaded with dexamethasone (DEX-NP) or vehicle (CON-NP) twice per week using a 30-gauge needle with a Hamilton glass microsyringe (50-μL volume; Hamilton Company, Reno, NV, USA). After withdrawing the needle, Neomycin plus Polymyxin B Sulfate antibiotic ointment (Sandoz, Princeton, NJ, USA) was applied to the eyes, and mice recovered on a warm pad. IOP was measured weekly using calibrated rebound tonometry (iCare TONOLAB, Vantaa, Finland). Both eyes were measured, and each recorded IOP value was an average of 6 measurements in each eye.

After 4 weeks of treatment with nanoparticles, all mice were bilaterally perfused *in vivo* with a 1:30,000 dilution of fluorescent, carboxylate-modified 100 nm FluoSpheres® (515 nm emission, Molecular Probes, Eugene, OR, USA) in PBS at a constant pressure of 15 mmHg for 1 hour, with the perfusion needle inserted in the anterior chamber. Mice were then sacrificed using isoflurane, and eyes were carefully enucleated, kept in cold PBS, then shipped to Georgia Institute of Technology overnight (Figure 1A). Following enucleation, a small incision was made in the superior quadrant of the globe to serve as an orientation mark during subsequent processing. Eyes from four of the mice were also fixed by immersion in 4% paraformaldehyde (Thermofisher) overnight before they were shipped on dry ice. Of the 14 mice (28 eyes) used to collect IOP data, 19 eyes were successfully perfused and analyzed, while nine eyes (four DEX, five CON) were excluded due to technical issues with the tracer perfusion (Table 1). Detailed information on each eye used in this study is available in Supplemental Table 1.

**Figure 1:**
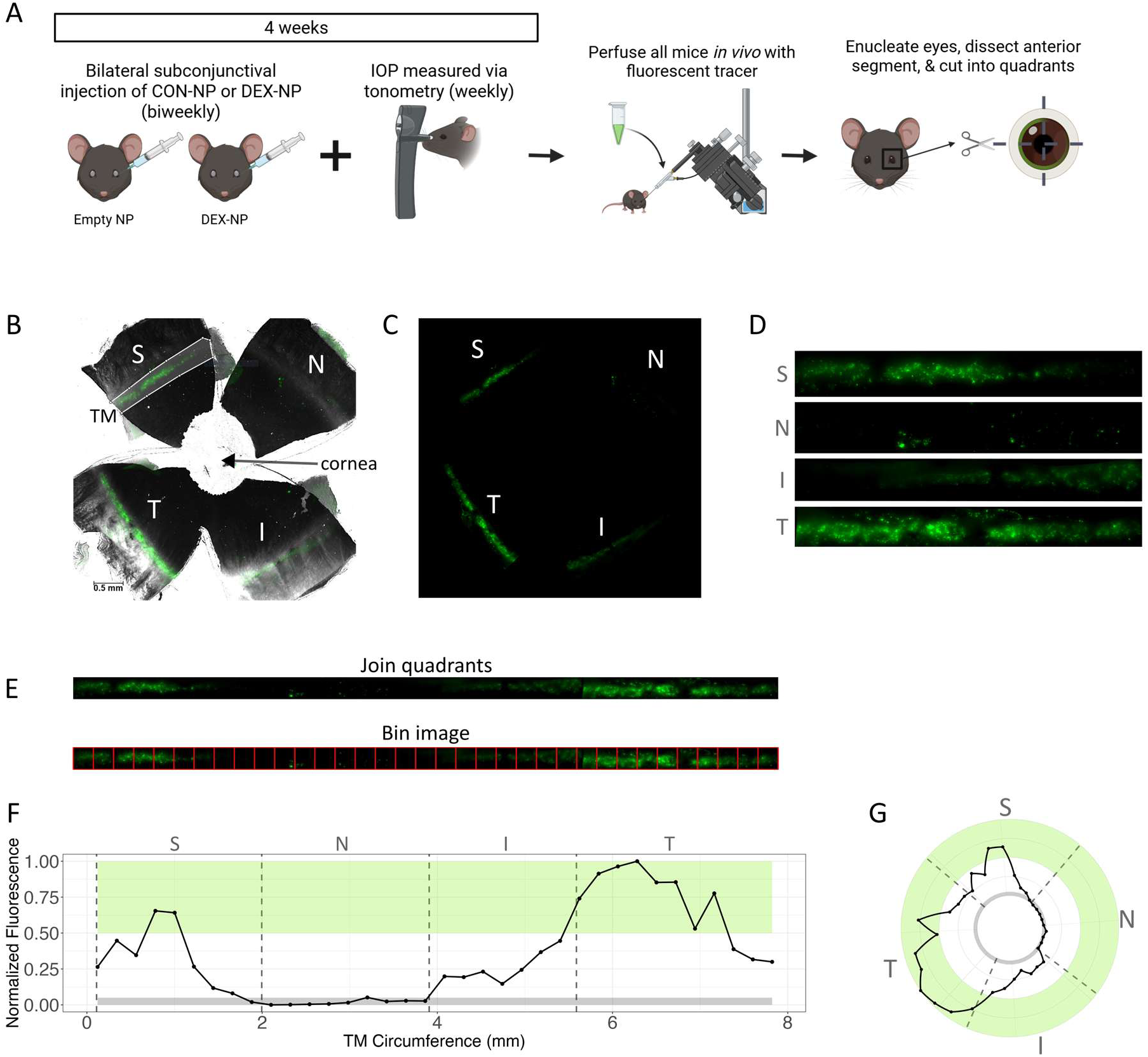
DEX-NP treatment and segmental flow quantification. **A)** Schematic of workflow for inducing ocular hypertension with DEX-NP treatment. Wild-type C57Bl/6 mice received bilateral DEX-NP or CON-NP delivery on a biweekly basis, and IOP was measured weekly via rebound tonometry. After 4 weeks of treatment, mice were perfused *in vivo* with Fluospheres before being euthanized, and their eyes were enucleated. Eyes were dissected to isolate anterior segments and then divided into quadrants. **B)** Raw image of a representative eye divided into quadrants after anterior segment dissection. Merged brightfield and fluorescence channels show the fluorescent tracer distributed around the limbus. The white outline represents our manual selection of the limbus for one quadrant. **C)** Masked fluorescence image of the eye after performing background correction and removing data outside of the selected limbus region. **D)** Straightened images of limbus quadrants. **E)** Straightened quadrant images were concatenated to represent the entire circumference in one strip, then divided into bins of 220 µm (red lines). **F)** Plot of mean fluorescence in each bin, normalized so values range between 0 and 1 to represent relative tracer distribution around the circumference of each eye. The light green region (top 50%) represents high flow, gray (bottom 5%) represents low flow, and values in between are taken as intermediate flow. Dotted lines represent quadrant boundaries. **G)** The data shown in (F) in a polar plot, matching the eye’s anatomy and allowing easy registration of quadrant locations. (*S=Superior, N=Nasal, I=Inferior, T=Temporal*)

**Table 1:** Summary of eyes used across experiments.

| Experiment | Number of Eyes Contributing Data to this Study |  |  |
| --- | --- | --- | --- |
|  | DEX eyes | CON eyes | Total (DEX + CON) |
| IOP measurement | 14 | 14 | 28 |
| Tracer perfusion | 10 | 9 | 19* |

\*9 eyes were excluded due to technical issues with tracer perfusion.

### 2.4. Tissue processing

After receipt of eyes at Georgia Tech, the anterior segment of each eye was dissected into four quadrants (superior, nasal, inferior, temporal), and each quadrant was imaged *en face* using a Leica DM6 microscope. Quadrants were positioned in a flat-mount configuration such that the fluorescent signal from the tracer in the TM could be imaged through the corneoscleral tissue. Both brightfield and fluorescent images were acquired at 10× magnification, and a GFP filter cube with the light source at full power was used for fluorescence imaging. The brightfield images were obtained using external overhead illumination using flexible gooseneck LED lights to penetrate the corneoscleral region.

Based on the *en face* fluorescence distribution within each eye, we identified TM regions with the highest fluorescent signal (HF) and lowest fluorescent signal (LF). We then collected sagittal cryosections from the HF and LF regions of interest for stiffness measurements and immunofluorescence labeling. To ensure that we could reliably collect cryosections from a pre-selected region of a quadrant, we performed preliminary calibrations with the microscope and cryostat, as described in the supplemental materials (Supplemental Figure 1 & Table 2).

After verifying the agreement between LASX software annotations and cryostat length measurements, we used a similar technique on the quadrants containing HF and LF regions to collect sagittal sections. Quadrants were embedded in optimal cutting temperature compound (OCT), snap frozen in 2-methylbutane cooled with liquid nitrogen, and stored at −80 °C. The distance of each HF and LF region from the quadrant edge was annotated in the Leica DM6 LASX software, and those distances were used to accurately collect serial cryosections from both HF and LF regions using the Cryostar NX70. When possible, regions where high and low flow areas occurred in close proximity were prioritized, as it was judged that these were more likely to reflect true biological differences rather than artifacts introduced by placement of the needle used to infuse the tracer.

### 2.5. Segmental Flow Analysis

Tracer distribution analysis methods were adapted from Reina-Torres et al.^29^. Image processing was performed in MATLAB (R2023b, Mathworks, Natick, MA). Using the overlay of the brightfield and fluorescent images, the limbus was manually outlined using the polygon tool guided by anatomical landmarks and fluorescent signal from the tracer. The outlined regions of interest were saved as masks (Figure 1B). To account for background fluorescence, three regions of tissue without tracer (one from the cornea and two from the sclera) were manually selected, and the mean pixel intensity of these regions was subtracted from the fluorescent channel image. Quadrant images were masked based on the previously drawn outlines of the limbus in each quadrant to isolate the relevant fluorescent signal (Figure 1C), and each quadrant’s masked limbus image was straightened (Figure 1D). Images were then concatenated to form one image to represent the entire circumference of the eye, and the resulting image was split into 220 mm wide bins (Figure 1E). Within each bin, the mean pixel intensity was calculated, and values were normalized within each eye by scaling the minimum to 0 and the maximum to 1 to account for inter-sample variability in total fluorescence intensity. Bins with normalized fluorescence greater than 0.5 were taken as high flow, those with normalized fluorescence less than 0.05 were taken as low flow, and bins in between were taken as intermediate flow, based on methodologies used in existing segmental flow literature, where 50% of the maximum fluorescence was taken as high flow^22,30,31^ (Figure 1F). To visualize fluorescence values registered by quadrant around the circumference of the eye, data was also displayed using polar plots (Figure 1G).

### 2.6. Atomic Force Microscopy

Sagittal 16 µm thick anterior segment sections from HF and LF regions of each eye were cut on a CryoStar NX70 cryostat (Thermofisher). Sections were placed on Superfrost Plus Gold slides (Fisher), allowed to dry, and stored at −80 °C. Prior to AFM measurements, the samples were thawed and the OCT washed away by submerging the slide in PBS for at least 10 min at 4 °C. All AFM measurements were conducted within one week of sectioning and slides were submerged in room temperature PBS during AFM measurements (1–4 hours) to maintain tissue hydration.

An MFP-3D AFM (Asylum Research, Santa Barbara, CA) with a prefabricated 10 µm diameter spherical borosilicate glass probe (Novascan, Boone, IA) attached to a V-shaped silicon nitride cantilever (nominal spring constant 0.1 N/m) was calibrated using the thermal noise method ^32^ and used to obtain a 4 *x* 4 raster scan (i.e., a force map) of measurements in the sclera, cornea, and TM. Each force map covered a 15 *x* 15 µm area (Figure 2). For each measurement, the probe approach velocity was 1 µm/s, probe retract velocity was 5 µm/s, x-y velocity during force mapping was 1 µm/s, and the trigger force was 7 nN. Each force map was immediately repeated in the same location, and the effective Young’s modulus was averaged between the two measurements at each measurement location in the force map to estimate the stiffness at each location.

**Figure 2:**
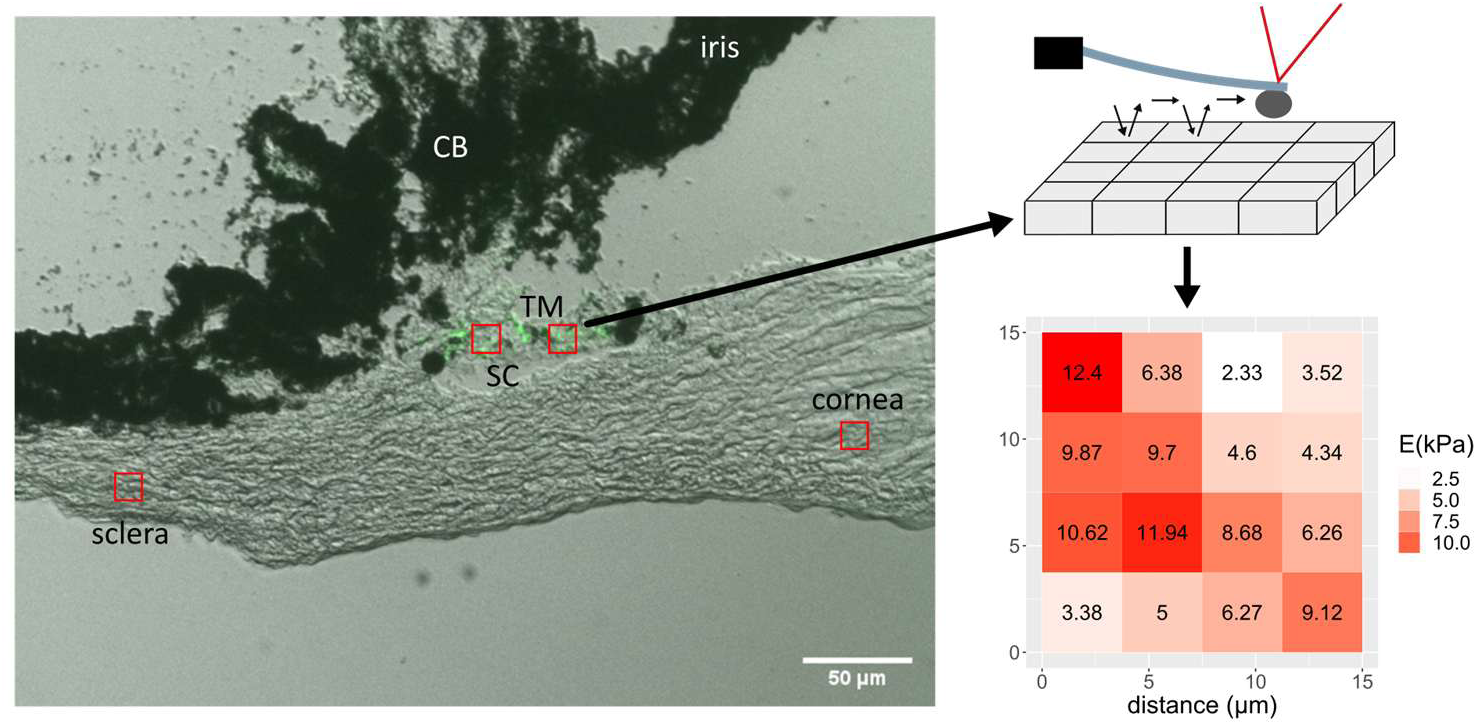
Schematic of AFM force mapping. Representative anterior segment cryosection with fluorescent overlay shows Fluospheres (green) present in TM. Red boxes indicate locations where 15 × 15 µm force maps were collected in the sclera, cornea, and TM. Each force map contains a 4 × 4 grid of measurements, so the effective Young’s modulus can be calculated across the boxed region as shown.

### 2.7. AFM Data Analysis

Force mapping data was analyzed as previously described^33^. In brief, the Hertz model for a spherical indenter was used to fit all force-displacement curves and thereby determine the effective Young’s modulus at each location using the following formula: 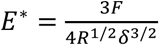, where *R* = probe radius, 𝛿 = indentation depth, and *F* = applied force. Curve fitting was performed using a custom R script, and the full indentation depth was used for curve fitting.

After fitting force-indentation curves according to the Hertz model, each curve fit was manually evaluated. Curves with a poor fit to the Hertz model, defined as having a residual greater than 50 nN, were removed from analysis (Figure 3A). Curves with an indentation depth greater than 2 µm were also removed from the analysis to avoid overestimating apparent Young’s modulus values due to substrate effects^34^ (Figure 3B). Additionally, if one or both force curves taken at the same location were removed due to a poor Hertzian model fit or large indentation depth, that measurement location was entirely removed from the analysis.

**Figure 3:**
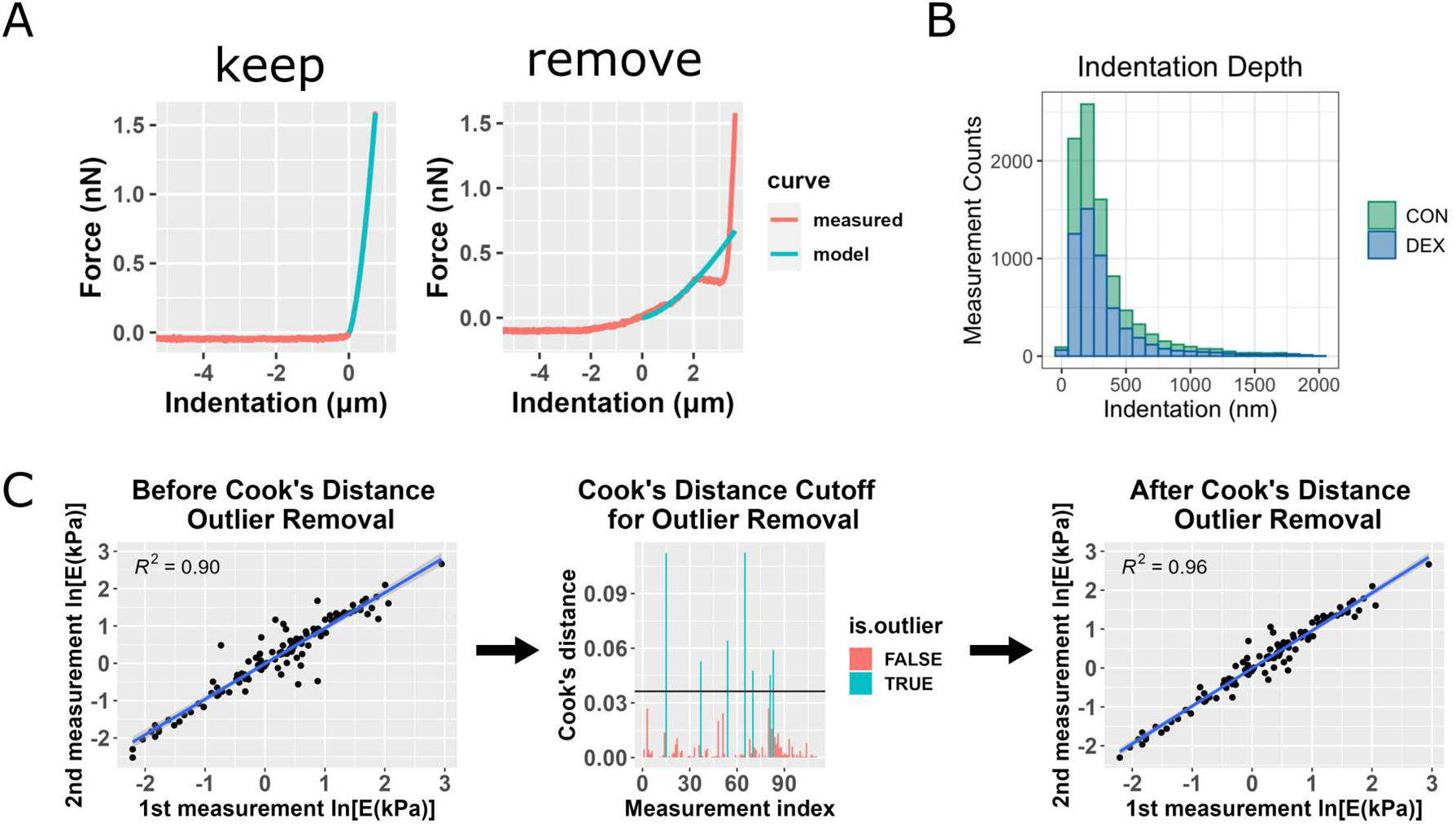
AFM data filtering and Cook’s outlier removal. **A)** Representative force-indentation plots (red) illustrating curve fitting quality, with the Hertz model fit shown in blue. The “good” fit (left) demonstrates a reliable curve that would be retained for further analysis, while the “poor” fit (right) exhibits inadequate fitting and would be excluded from the analysis. **B)** Histogram of all measurement indentation depths, colored by NP-treatment, including cornea, sclera, and TM data. Most measurements did not exceed a 1μm indentation depth, and any measurements with an indentation depth greater than 2 μm were removed from the analysis. **C)** Overview of the use of Cook’s distance outlier removal. Log-transformed effective Young’s modulus estimates from the first and second measurements at the same location for one eye are plotted against each other and linearly regressed (left plot). Cook’s distance is used to determine outliers (middle plot, 7 outliers shown in blue), indicating discordance between repeated measurements at the same point, and the regression is re-plotted without outliers (right plot). This process was applied to data from each eye.

By repeating each force map at each measurement location, we were able to use a test-retest paradigm to verify Young’s modulus measurements. Specifically, agreement between the two measurements at the same location provided a criterion to confirm repeatability of the measurements. For each eye, the fitted Young’s modulus from the second measurement was linearly regressed on the fitted Young’s modulus from the first measurement, for all measurement locations. Cook’s distance was calculated for each data point, and measurement locations for which the Cook’s distance exceeded the cutoff *4/N*, where N = number of data points ^35^, were removed from the analysis (Figure 3C).

### 2.8. Immunofluorescence Staining

Sagittal 10 µm thick anterior segment sections from HF and LF regions of each eye were collected with a CryoStar NX70 cryostat (Thermofisher) and fixed in 4% paraformaldehyde, followed by permeabilization in 0.2% Triton X-100 for 5 minutes and blocking in 10% goat serum for 30 minutes at room temperature.

Sections were then incubated overnight at 4 °C with primary rabbit monoclonal antibody against fibronectin (Abcam, ab2413; 1:500 dilution) and mouse monoclonal primary antibody against alpha-smooth muscle actin (α-SMA) conjugated to Cy3 (Sigma, C6198; 1:400 dilution), followed by overnight incubation at 4 °C with Alexa Fluor 647 goat anti-rabbit secondary antibody to visualize the fibronectin (FN) primary (Invitrogen, A-21245; 1:200 dilution). Each slide also included 1-2 negative control sections prepared in the same way, except that the primary antibody was omitted. Nuclei were counterstained using NucBlue Live Cell Stain ReadyProbes (Invitrogen, Waltham, MA, USA) according to the manufacturer’s instructions. Glass coverslips were mounted with Prolong Gold Antifade mounting medium (Invitrogen, Waltham, MA, USA).

Images were acquired on a Leica DM6 epifluorescence microscope equipped with a 40× objective lens and an additional 1.6× intermediate magnifier lens (total magnification: 64×). Identical exposure times and gain settings were used for all samples, and extended depth of field images were saved from z-stacks at 0.99 pixel/μm resolution. Fluorescence signals were detected using the following filter cubes: GFP filter cube (for the flow tracer), DSR filter cube (for α-SMA), A4 filter cube (for nuclei), and Y5 filter cube (for FN).

### 2.9. Immunofluorescence Quantification

Quantification of fluorescent signal was performed on 5-10 sections per HF and LF region of each eye using ImageJ (Fiji)^36^. To quantify FN and α-SMA intensities, background correction was performed by subtracting the mean fluorescence intensity from the sclera of each section, since we did not expect differences in scleral FN and α-SMA levels between high vs. low flow regions. A region of interest (ROI) was manually drawn around the TM using the polygon selection tool based on anatomical features from the brightfield channel (termination of Descemet’s membrane and SC lumen) and the location of fluorescent tracer visible in the GFP channel. We then calculated the mean gray value of both FN and α-SMA within the TM ROI.

To quantify the signal from green flow tracer, the green (GFP) channel was isolated in ImageJ, and the integrated density (defined as the total fluorescence intensity within a selected area) within the previously drawn TM ROI was measured. To correct for tissue autofluorescence and other nonspecific background signal, a second ROI was drawn in a nearby background region lacking beads, and its mean intensity was measured. A background-corrected integrated density (𝐼𝑛𝑡𝐷𝑒𝑛) was then calculated as: 𝐶𝑜𝑟𝑟𝑒𝑐𝑡𝑒𝑑 𝐼𝑛𝑡𝐷𝑒𝑛 = 𝐼𝑛𝑡𝐷𝑒𝑛_ROI_ − (𝐵𝑎𝑐𝑘𝑔𝑟𝑜𝑢𝑛𝑑 𝑀𝑒𝑎𝑛 × 𝑅𝑂𝐼 𝑎𝑟𝑒𝑎). This approach yields the total bead signal per ROI, normalized for differences in background fluorescence across images. However, because some sections had high background fluorescence, some of the resulting 𝐼𝑛𝑡𝐷𝑒𝑛 values were negative.

### 2.10. Statistical Analysis

#### 2.10.1. Intraocular Pressure

All statistical analyses were performed in RStudio (R version 4.2.0) using the “rstatix” and “fitdistrplus” packages. IOP values were averaged between the two eyes of each mouse, and each mouse was treated as a biological replicate. Outliers were identified and removed for each time point, and IOP data were analyzed using one-way ANOVA, followed by pairwise two-sided t-tests with Bonferroni correction for multiple comparisons.

#### 2.10.2. Flow Tracer Distribution

Because fluorescence values were normalized on a per-eye basis to range from 0-1, left and right eyes from the same animal were treated as independent samples for all downstream segmental flow analyses. Thus, the data reflect relative flow dynamics unique to that eye rather than systemic mouse-specific effects.

To compare distributions of normalized fluorescence intensity values between groups, a two-sample Kolmogorov-Smirnov (K-S) test was performed. This non-parametric test assesses whether two distributions differ in their cumulative distribution functions and does not rely on summary statistics such as the mean or underlying normality, making it appropriate for the normalized data in a defined range. In addition, kernel density estimation (KDE) plots were used as a non-parametric method to estimate the probability density function of the data. Both histograms and KDE visualization were used to illustrate the shape of the fluorescence intensity distributions and highlight regions where differences between groups were most pronounced. All other results are reported as mean ± standard deviation unless otherwise specified. Percentages of high, intermediate, and low flow were compared using two-sample t-tests with Bonferroni adjustment for multiple comparisons. This approach was justified because each percentage was derived from > 30 bins per sample (typically 38-40), allowing treatment of the percentages as continuous variables, and the Shapiro-Wilk test confirmed that the percentage values were approximately normally distributed. Statistical significance was defined as p < 0.05.

#### 2.10.3. Atomic Force Microscopy

Based on the AFM data from each tissue of each eye, we created histograms of Young’s modulus values after outlier removal, and a log-normal distribution was fit to Young’s modulus values using the “fitdistrplus” package in RStudio ^37^. We confirmed the log-normal distribution with a Kolmogorov-Smirnov (K-S) test, where a critical p-value of 0.05 was used. Then, we log-transformed the data and repeated the K-S test to confirm normally distributed data. We also visualized Q-Q (quantile-quantile), CDF (cumulative distribution function), and P-P (probability-probability) plots to verify that the normal distribution was a good fit to the log-transformed data (Figure 4). Based on the mean of the fitted normal distribution, we calculated a multiplicative (geometric) mean and multiplicative standard deviation to characterize Young’s modulus values in the non-transformed domain^38^. Detailed validation of this analysis pipeline has been previously reported^33^.

**Figure 4:**
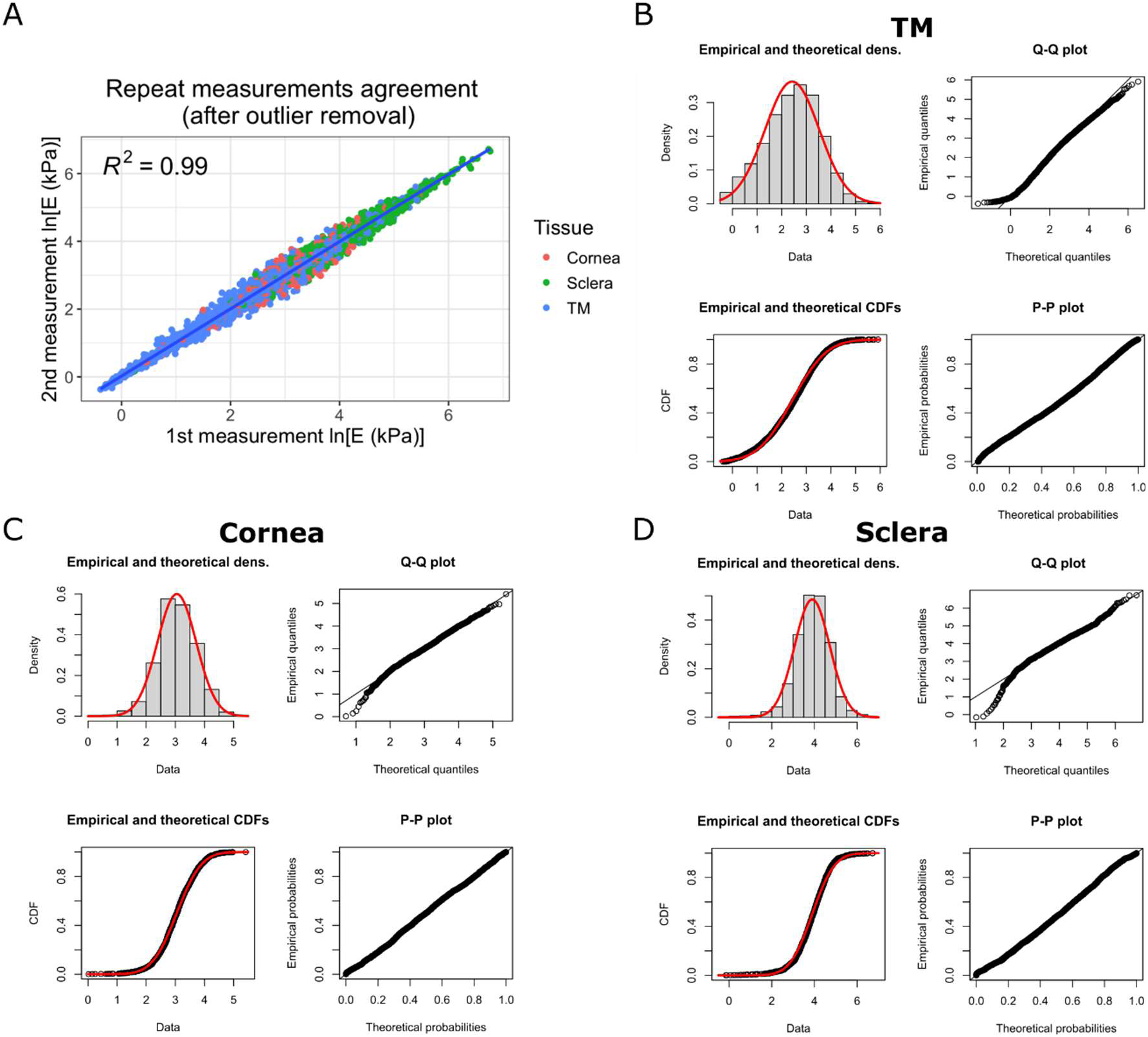
Test-retest agreement and log-transformation of Young’s modulus values. **A)** Linear regression of log-transformed effective Young’s modulus estimates from the first and second measurements at the same location for all cornea, sclera, and TM measurements. After data processing and outlier removal, there was strong agreement between repeated measurements at the same location (R^2^ = 0.99). **B-D)** The log-transformed Young’s modulus values from TM (B), cornea (C), and sclera (D) measurements appeared to be well-fit by a normal distribution, as judged by a histogram of Young’s modulus values vs. a fitted normal distribution (top left), and by comparisons of actual and theoretical quantiles (top right), actual and theoretical cumulative distribution functions (bottom left), and actual and theoretical probability distributions (bottom right). In all four panels, actual data is in black/grey and theoretical fits are overlain in red.

Group differences in geometric means were compared using ANOVA, followed by unpaired t-tests with Bonferroni correction for pairwise comparisons. For within-eye comparisons between HF and LF regions, pairwise t-tests were used.

#### 2.10.4. Immunofluorescence

To compare mean FN and α-SMA intensity levels in the TM between DEX-and CON-treated mice, mean fluorescence intensity values were analyzed using a linear mixed-effects model with each mouse treated as a random effect, and each section as a repeated technical measurement nested within each mouse. Only one eye from each mouse was used. Treatment (DEX-NP vs. CON-NP) and flow region (HF vs. LF), and their interaction were treated as fixed effects. Model fitting was performed in R using the lme4 package. For visualization, model-estimated means and individual measurements were normalized to the mean of the CON–HF group, and error bars represent 95% confidence intervals.

Linear regression analysis was also used to assess the relationship between the fluorescent tracer levels vs. FN and α-SMA labeling. For each sample and nanoparticle treatment condition, mean FN or α-SMA fluorescence intensity was regressed against tracer background-corrected integrated density (𝐼𝑛𝑡𝐷𝑒𝑛). Extreme outliers for integrated density were removed. From each regression model, the slope, intercept, coefficient of determination (R²), and p-value for the tracer fluorescence term were extracted. P-values were adjusted for multiple comparisons using Bonferroni correction. Regression models and results were visualized by plotting FN or α-SMA mean intensity as a function of tracer fluorescence, with best-fit regression lines overlaid for each sample and treatment condition.

## 3. Results

### 3.1. Intraocular Pressure

At baseline, the control mouse cohort exhibited slightly higher baseline IOP compared to the DEX-treated cohort (20.0 ± 0.44 mmHg vs. 19.1 ± 0.72 mmHg, p = 0.039). Mice receiving DEX-NPs showed significantly elevated IOP after one week of treatment (p < 0.0001), and this difference persisted over the course of four weeks (Figure 5A). On average, DEX treatment increased IOP by 6.35 ± 1.85 mmHg (33.3%) from baseline to week 4 (p < 0.0001), whereas animals injected with empty NPs exhibited a modest, non-significant decrease of 0.45 ± 0.87 mmHg (2.19%) (Supplemental Figure 2). Considering the higher baseline IOP values in the control mice, this result further emphasizes that the DEX-NP treatment causes ocular hypertension. At week 4, DEX-treated eyes also displayed greater variability in IOP compared to controls (standard deviation = 2.77 vs 1.17 mmHg), suggesting some heterogeneity in the number of DEX-NPs injected and/or magnitude of the response to DEX-NPs.

**Figure 5:**
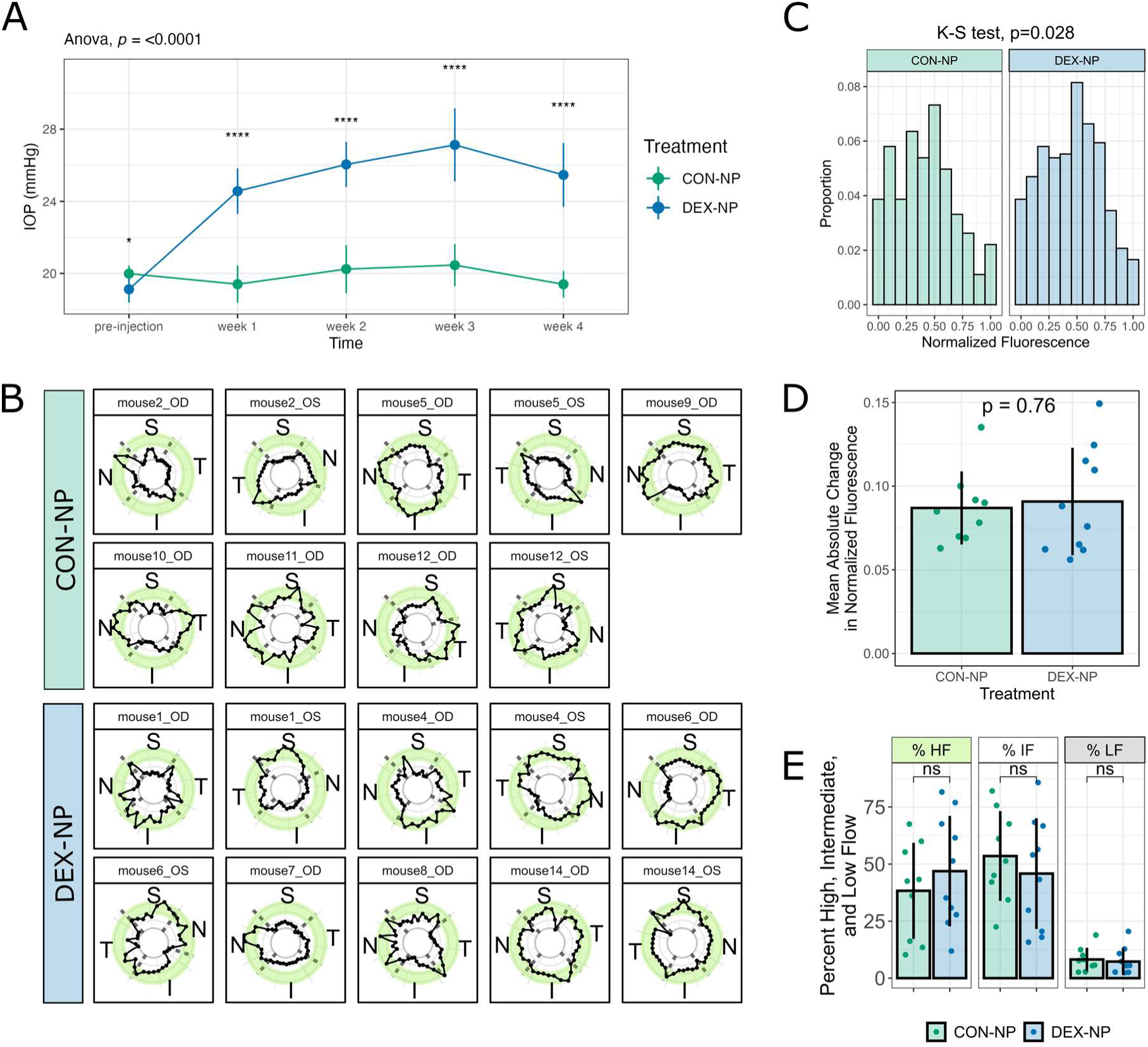
DEX-NPs cause ocular hypertension and impact segmental flow distribution. **A)** IOP of DEX-NP over the course of the treatment was significantly elevated compared to CON-NP-treated mice. Data is shown as mean ± standard deviation. (p < 0.001 by ANOVA; pairwise comparisons are based on t-test with Bonferroni-adjustment, **** p < 0.001). **B)** Normalized fluorescence intensity values around the circumference of all analyzed DEX-NP and CON-NP-treated eyes. Points in the shaded green area are considered HF, and values in the gray shaded area are considered LF. Sample names identify eyes from the same mouse. **C)** Histograms of binned normalized fluorescence values from all DEX- and CON-treated eyes. The two distributions are significantly different (K-S test, p=0.028). **D)** Mean absolute change in normalized fluorescence between adjacent data points (bins) shows no difference between cohorts. Individual points represent each eye, and error bars show standard deviation. Group means were compared with an unpaired t-test. **E)** Percentage of HF, IF, and LF in DEX-treated eyes vs. Controls. Each point represents an individual eye, and error bars show standard deviation. Mean percentage values were compared using an unpaired t-test with Bonferroni adjustment.

### 3.2. Segmental Flow Distribution

We observed a considerable amount of variability in tracer distribution between animals, and even between eyes from the same animal (Figure 5B). We did not see any trends in terms of specific anatomic quadrants having more outflow than others. When comparing the distribution of normalized mean fluorescence values as a histogram, we observed significant differences between CON-NP and DEX-NP-treated eyes, as indicated by a Kolmogorov-Smirnov test (Figure 5C, K-S test p=0.028). The kernel density estimates suggest that the difference is concentrated in the intermediate flow region, towards the center of the distribution (Supplemental Figure 3). The control group exhibited greater density in the lower-mid range of normalized fluorescence values (approximately 0.2-0.4), whereas the DEX-treated group had relatively higher density in the upper-middle range of values (approximately 0.5-0.6). Skewness values (0.05 in the DEX-treated group, 0.31 in the control group) indicate that both distributions were approximately symmetric, but the CON-treated values were slightly more right-skewed.

To capture local variation in fluorescence intensity around the limbus, we calculated the mean absolute difference between adjacent normalized intensity values. This measure reflects the average change in signal from one bin to the next, with greater values indicating greater spatial heterogeneity in tracer distribution. Using this approach, we found no significant difference between cohorts (Figure 5D), indicating no differences in the variability in fluorescence distribution around the circumference of the eye with DEX-NP treatment. We further quantified the percentage of high, intermediate, and low flow in each eye, and did not observe any significant differences between the DEX-treated eyes vs. controls (Figure 5E). Eyes were treated as independent samples, as there was no correlation in percentage of high, intermediate, or low flow between contralateral eyes (Supplemental Figure 4). This result suggests that DEX-induced ocular hypertension did not shift the overall average outcome of high, intermediate, and low flow regions, or the variability in relative outflow around the circumference of the eye, but DEX-treatment resulted in a shift in distribution within the intermediate flow range values.

### 3.3. Cornea, Sclera, and Trabecular Meshwork Stiffness

We first confirmed differences in stiffness between cornea, sclera, and TM tissues (Figure 6). In both dexamethasone-treated and control eyes, the sclera exhibited the highest stiffness, followed by the cornea and then the TM. These findings are consistent with prior reports demonstrating that the sclera is stiffer than the cornea^39^, and align with data previously reported by our group in rats^33^. Together, these results validate that the force mapping approach employed here can detect physiologically relevant differences in ocular tissue stiffness.

**Figure 6:**
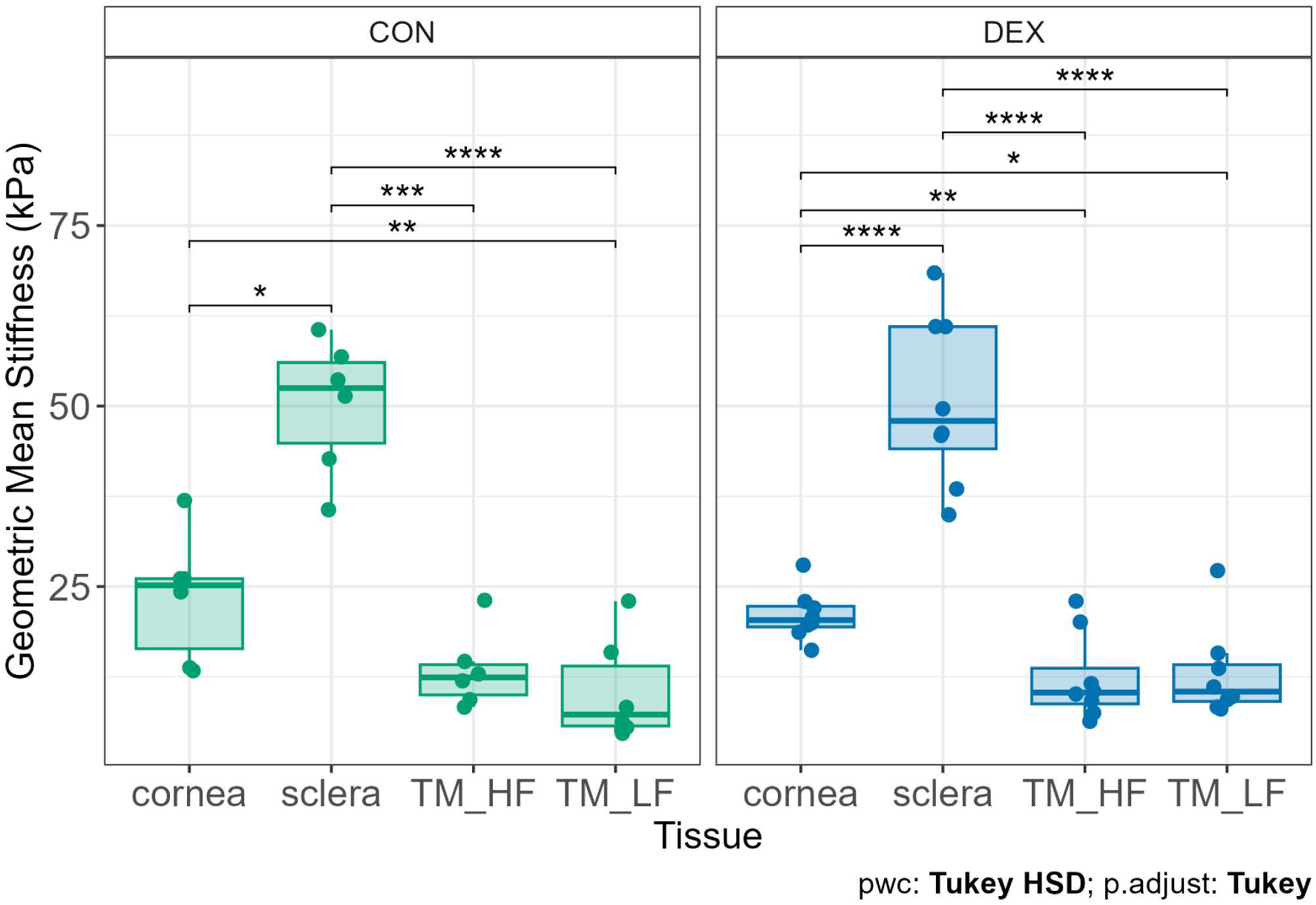
Stiffness of cornea, sclera, and TM in CON and DEX mice. In both treatment groups, the sclera was significantly stiffer than both the cornea and TM, and the cornea was stiffer than the LF TM regions. In the DEX-treated group, the cornea was also significantly stiffer than HF TM regions. Individual points represent geometric mean stiffness values from each eye. Statistics were performed using a one-way ANOVA followed by Tukey’s HSD for pairwise comparisons (*p<0.05, **p<0.01, ***p<0.001, ****p<0.0001).

### 3.4. TM stiffness in DEX vs. CON and HF vs. LF regions

Next, we directly compared ocular tissue stiffness in DEX-NP and CON-NP-treated eyes. Despite differences in IOP, no differences were detected in the geometric mean stiffness of the TM as measured by force mapping (Figure 7A). We also did not observe any differences in cornea or sclera stiffness between these groups. Further, a pairwise comparison of HF and LF regions from each eye showed no significant differences in stiffness between segmental flow regions in both CON-NP and DEX-NP eyes (Figure 7B).

**Figure 7:**
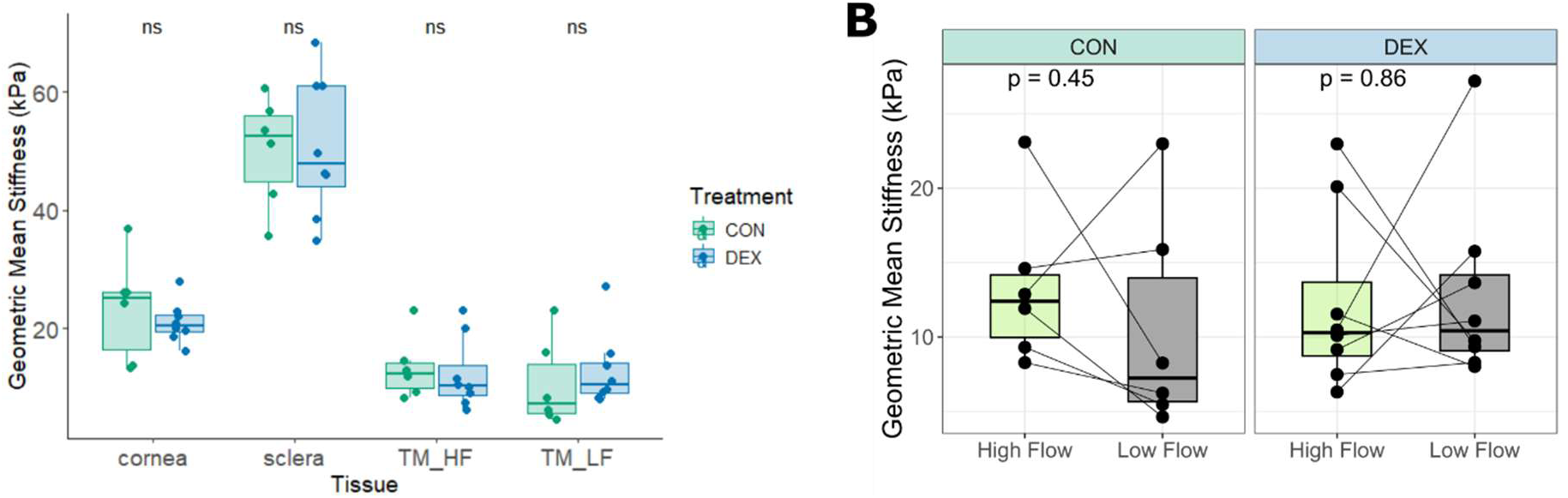
DEX vs. CON and HF vs. LF TM stiffness. **A)** Comparison of DEX-NP and CON-NP treated cornea, sclera, and high and low flow TM stiffnesses. Each point represents the geometric mean of the effective Young’s modulus values from one eye. (t-test with Bonferroni adjustment for pairwise comparisons; ns=not significant) **B)** Paired t-test comparing high and low flow regions from the same eye reveals no significant difference between HF vs. LF TM stiffness in both CON-NP and DEX-NP-treated eyes. Lines connect HF and LF regions from the same eye.

We further analyzed the maximum and minimum modulus values extracted from each TM force map to assess the range of tissue stiffness within HF and LF regions. Maximum stiffness values did not differ significantly between DEX-NP and CON-NP eyes in either region. However, analysis of minimum stiffness values revealed a significant difference in LF regions (p = 0.0428), with CON-NP eyes exhibiting lower minimum stiffness compared to DEX-NP eyes (Figure 8). This finding suggests that, while overall TM stiffness was comparable between treatment groups, certain localized regions within the LF TM of control eyes were softer, potentially providing pathways that facilitate aqueous humor outflow.

**Figure 8:**
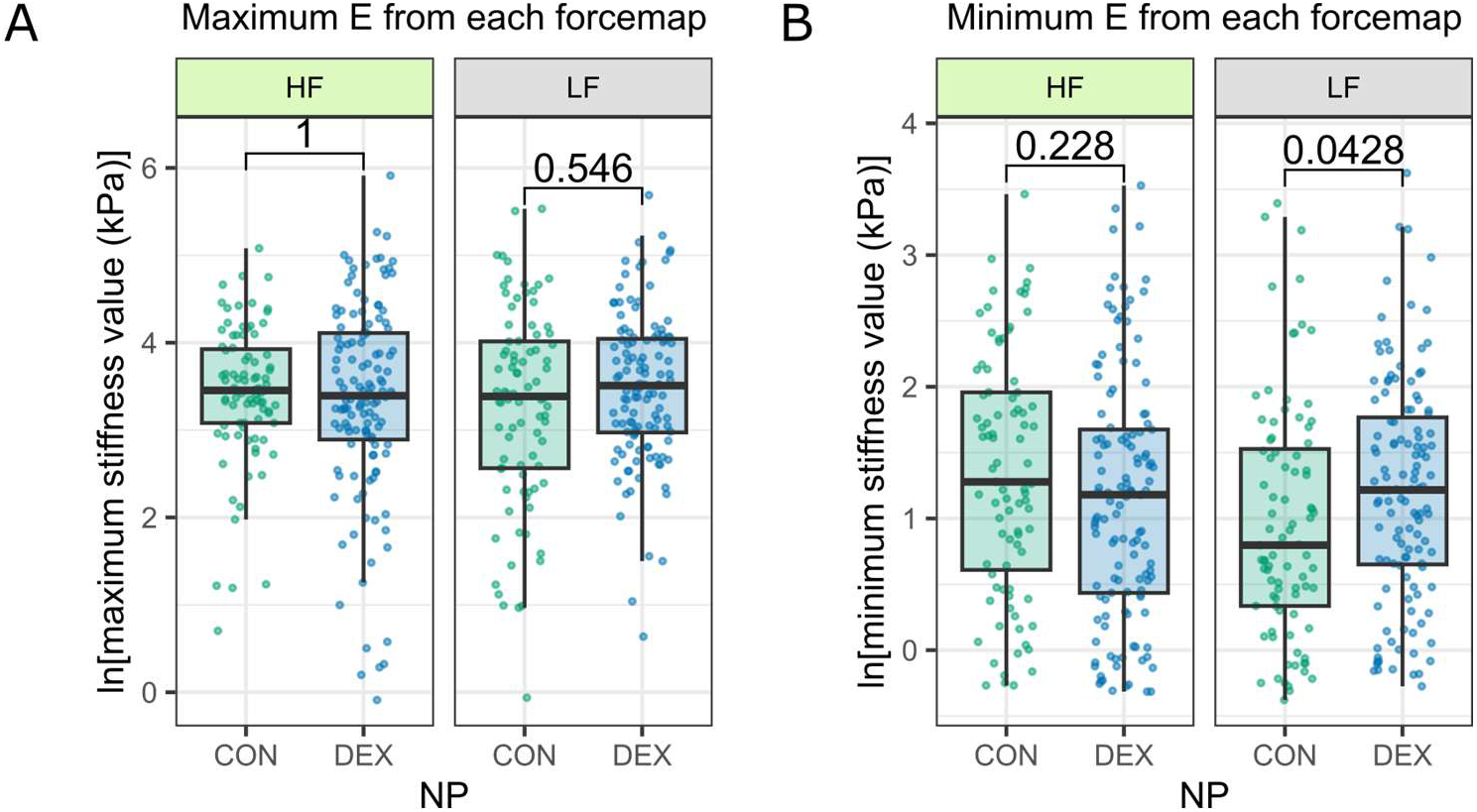
Maximum and minimum stiffness values from each force map. **A)** There were no significant differences in maximum stiffness values between DEX-NP and CON-NP-treated eyes in the high or low flow TM regions. **B)** When comparing minimum stiffness values from each force map, in the LF regions, the CON-NP eyes had lower minimum stiffness values. p-values shown are from Wilcoxon rank-sum tests with Bonferroni correction for multiple comparisons. E = Effective Young’s Modulus.

### 3.5. Fibronectin and α-SMA

To explore other differences in the TM between DEX-treated and CON-treated mouse eyes that might underlie the observed IOP difference, we investigated two major TM matrix and cytoskeletal components, FN and α-SMA, using immunofluorescent staining. FN is a major component of the TM ECM and is known to be upregulated by DEX in both TM cells and tissue^7,24,40,41^. Similarly, α-SMA levels have been shown to increase in DEX-treated mouse TM compared to controls^41,42^. On average, FN levels were greater in DEX-treated mice compared to controls (DEX: 1.33 ± 0.589 a.u. vs. CON: 0.484 ± 0.278 a.u.). Linear mixed-effects modeling revealed a significant interaction between treatment and flow region (p < 0.001), and post-hoc comparisons using model-adjusted means showed that DEX treatment was associated with increased FN levels in the LF regions compared with controls (1.19-fold vs CON-NP-treated HF, p < 0.001), whereas there was no significant difference in the HF region (Figure 9), consistent with previous reports that DEX induces FN upregulation^40,43,44^. We also investigated whether FN expression correlated with tracer intensity, but no consistent relationship was detected across groups (Figure 10). While one DEX-treated mouse (Mouse 8) demonstrated a significant negative correlation (p < 0.05), this pattern was not observed in the other DEX eyes or in any of the control eyes.

**Figure 9:**
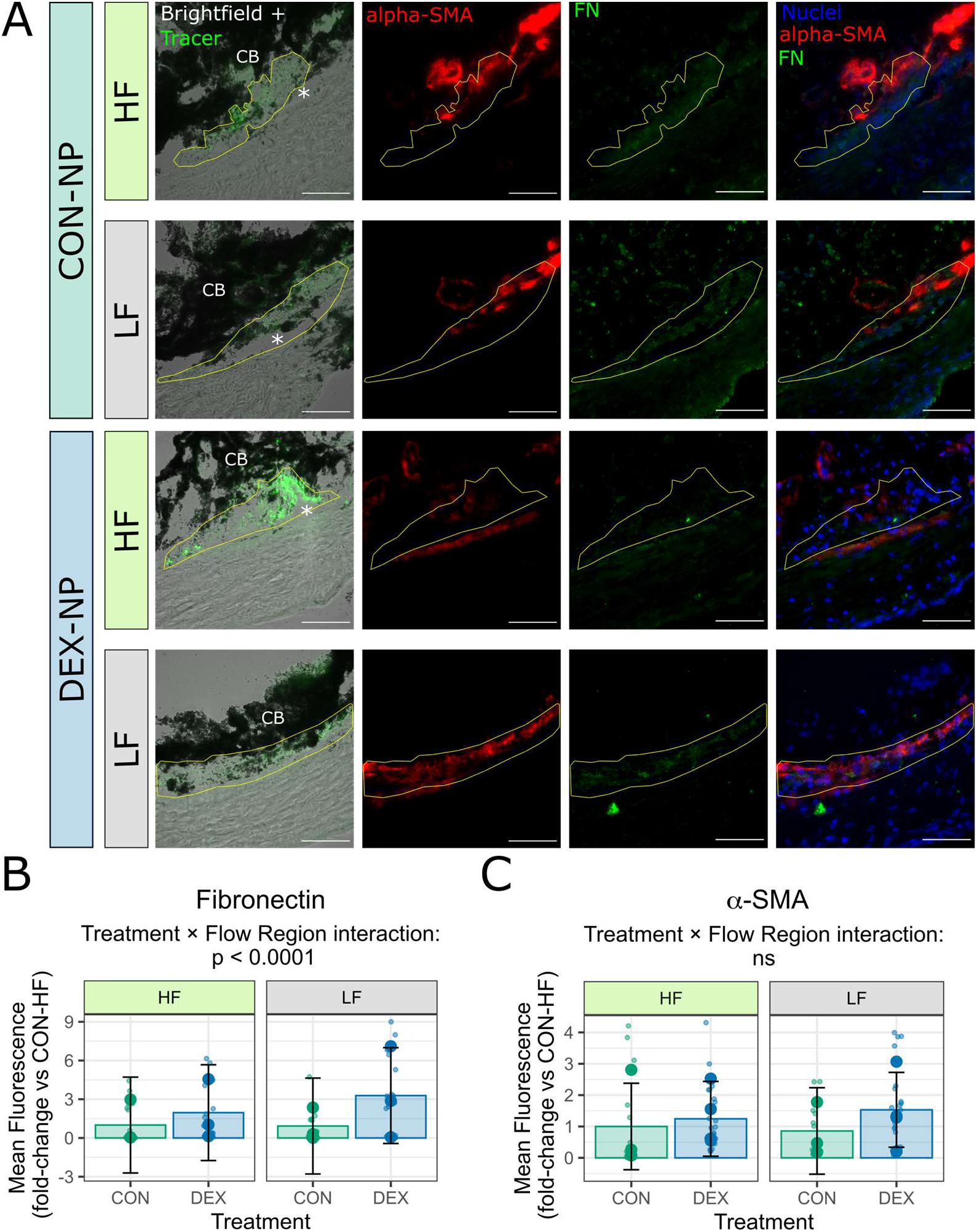
Fibronectin and α-SMA labeling in HF and LF regions of DEX-treated and CON-treated mice. **A)** Representative sections of immunofluorescent staining of FN and α-SMA in mouse trabecular meshwork cryosections. The first column of panels shows the overlay of the brightfield channel with the Fluospheres, visible in the GFP (green) channel. The following columns show α-SMA and FN, respectively, and the final column is an overlay of the α-SMA, FN, and nuclei staining. The TM is shown by the area outlined in yellow, and an asterisk (*) denotes the SC lumen, except in the final row, where the canal was collapsed. CB = ciliary body. Scale bar = 50 µm. **B)** Quantification of the mean FN labeling in DEX-treated and CON-treated eyes, split by flow region. Each large symbol represents the mean FN intensity from one eye, while each small symbol represents the mean FN intensity in the TM of a single cryosection Each mouse was considered to be an independent sample since only one eye per mouse was used, and sections were treated as technical replicates (n=6 eyes, 4-8 sections per eye). Bars represent the mean and 95% confidence interval. A linear mixed-effect model revealed a significant interaction between treatment and flow region, and post-hoc comparisons using model-adjusted means showed that DEX treatment significantly increased FN levels in the LF regions compared with controls (p < 0.001) but did not have a significant effect in the HF regions. **C)** Quantification of the mean α-SMA intensity in the TM of DEX vs CON eyes, subdivided by flow region. Each large symbol represents the mean α-SMA intensity from one eye, while each small symbol represents the mean FN in the TM of a single cryosection. Each mouse was considered an independent sample since only one eye per mouse was used, and sections were treated as technical replicates (n=7 eyes, 4-8 sections per eye). Bars represent the mean and 95% confidence interval. No significant differences were detected with linear mixed-effect modeling between any groups.

**Figure 10:**
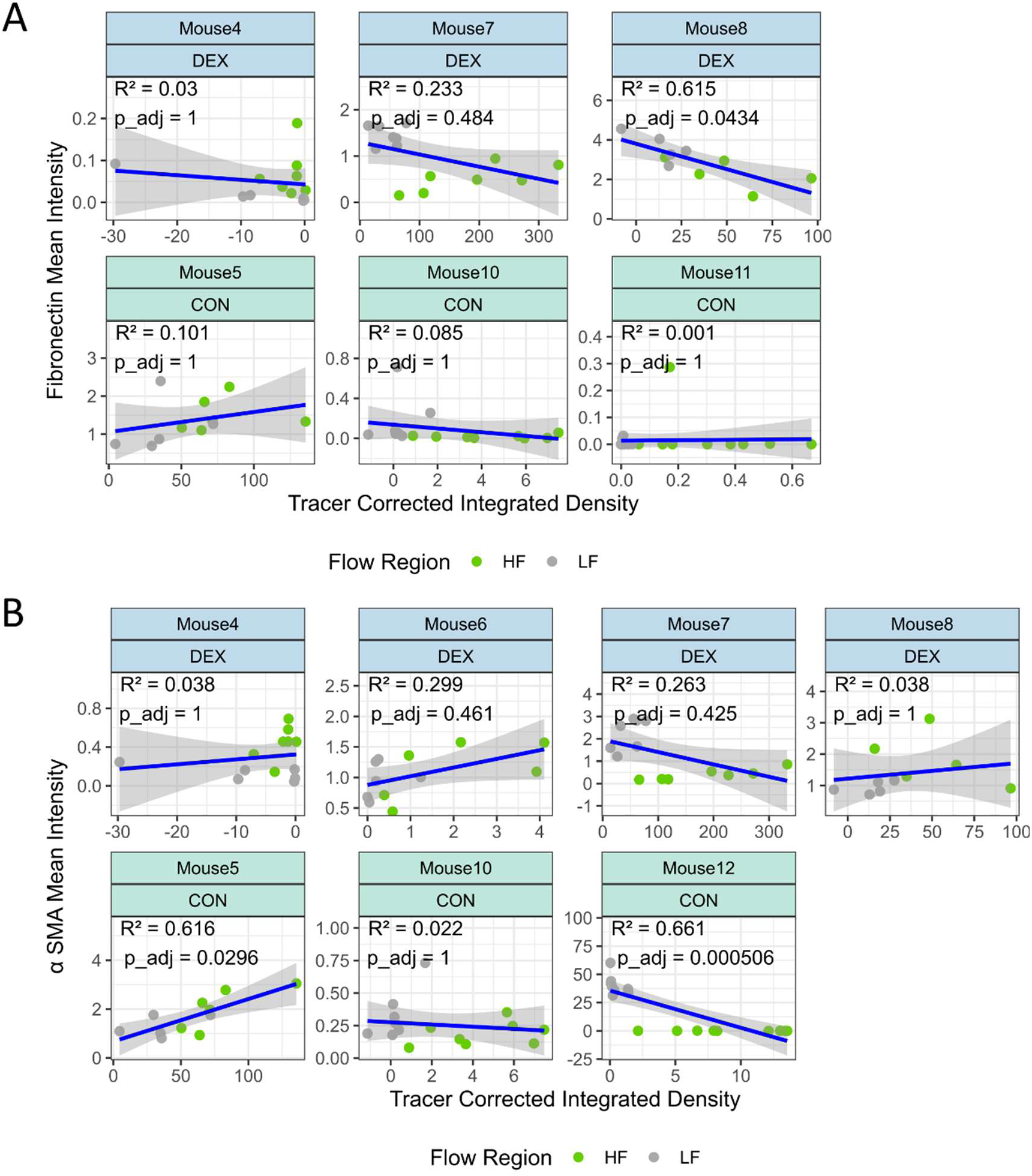
Correlation of fibronectin and alpha-SMA intensities with tracer intensity. Fibronectin (A) and α-SMA (B) mean integrated density plotted against the tracer corrected integrated density in DEX-treated and control mice. The blue line represents the linear regression fit and the gray shaded area indicates the 95% confidence interval, with coefficient of determination (R²) and p-value shown in the top left.

On average, α-SMA levels were higher in DEX-treated eyes compared to controls (DEX: 1.01 ± 0.253 a.u. vs. CON: 0.673 ± 0.330 a.u.) consistent with previous reports of DEX-induced α-SMA upregulation^41,42^. However, no significant differences were detected with linear mixed-effects modeling between LF and HF regions within either group, or in the interaction between DEX-treatment and flow region (Figure 9C). Correlation analysis between α-SMA and tracer intensity did not reveal consistent patterns (Figure 10B). Interestingly, two control eyes showed significant correlations, but in opposite directions—one positive and one negative—highlighting the variability in α-SMA labeling across sections. Prior work has also reported increased α-SMA labeling in the outer wall of SC following DEX treatment in mice^42^, and we observed a similar trend in some, but not all, of our samples as well (Figure 9A).

## 4. Discussion

To our knowledge, this is the first report of segmental flow quantification in an ocular hypertensive mouse model. While the mechanisms of steroid-induced glaucoma are not the same as those of POAG, they share many similarities, and thus these findings provide important insights for POAG research and disease models.

Based on existing literature on segmental flow in human donor eyes, we anticipated greater heterogeneity (segmentation) in ocular hypertensive DEX-treated eyes. Early work suggested “more segmentation” in glaucomatous eyes^12^, and anterior segment perfusion under elevated (2x) pressure also results in changes in segmental flow distribution^31^. Similarly, a study in ocular hypotensive SPARC-knockout mice reported disrupted outflow segmentation compared to controls^45^. However, SPARC is differentially expressed between high- and low-flow regions^18,22^, indicating a potential involvement in segmental outflow regulation which may underlie the observed changes in flow distribution in SPARC knockout mice that are not recapitulated in the dexamethasone model.

Interestingly, total raw fluorescence did not differ between control and DEX-treated eyes (Supplemental Figure 5), despite the expectation that the decreased outflow facility with DEX treatment would result in less total tracer present in the TM when delivered by a constant time and pressure perfusion. While unexpected, this finding underscores that total fluorescence in these eyes cannot be used as a proxy for total outflow. Instead, the relative distribution of tracer in the eye can be examined to gain insight into segmental flow patterns, justifying the normalization of fluorescence values within each eye used in the analysis.

Age is another important consideration. Previous work has shown that segmental flow patterns are highly dynamic in younger mice, with tracer distributions changing more drastically over the course of 14 days compared to the situation in older mice^29^. Since these experiments utilized young (2–4-month-old) mice, where the flow distribution around the eye was potentially quite dynamic, it is possible that the dynamic nature of the flow regions mitigated the effects of ocular hypertension on segmental flow regions, resulting in only subtle differences in between cohorts. Moreover, using young mice may have reduced the likelihood of detecting stable TM stiffness differences between HF and LF regions in the AFM studies. Future studies in older animals, as well as in human donor eyes, may help determine whether HF and LF regions exhibit consistent differences in tissue stiffness, particularly in the context of age-related ECM remodeling and primary open-angle glaucoma.

We were surprised to see that dexamethasone treatment did not alter the overall TM stiffness in our mouse model. Previous studies using AFM in rabbits have reported increased TM stiffness following treatment with dexamethasone eyedrops^7^; however, these animals did not exhibit elevated IOP. In mice, previous work using inverse finite element modeling based on *in vivo* OCT imaging also suggested increased TM stiffness after DEX-NP treatment^41^. Importantly, stiffness estimations are highly dependent on measurement methodology; when using OCT *in vivo*, the tensile response of the TM is being evaluated, while in AFM measurements on cryosections, the compressive response of tissue is evaluated. Removing AFM measurements with a large indentation depth to account for Hertz model assumptions may also result in exclusion of very soft regions of the tissue. Additionally, dexamethasone may also influence cell mechanics directly rather than impacting ECM stiffness, leading to differences between *in vitro*, *in vivo* and *ex vivo* studies. *In vitro* studies have shown that TM cells become stiffer following dexamethasone treatment, which is a response typically attributed to actin fiber formation, cytoskeletal remodeling, and altered cell–ECM interactions^7,24^. *In vivo*, DEX-NP-treatment has been reported to significantly increase IOP after just 3 days in mice; further, IOP elevation is attenuated by treatment with a rho-kinase inhibitor, which affects TM cell contractility, after just 4 days in mice. These rapid IOP changes are likely due to cellular mechanisms rather than large-scale changes in ECM deposition, which typically act on a slower time scale^11^. Such cellular-level changes likely contributed to the observed increased outflow resistance and rapid IOP elevation observed here, but their impact on tissue-level biomechanics may not be captured by bulk AFM measurements on cryosections, which primarily reflect ECM stiffness. Further, our findings are consistent with a previous AFM study on cryosectioned TM from DEX-treated mice that reported no significant TM stiffness differences^9^. We hypothesize that the bulk, compressive measurements employed here by using force mapping with a spherical probe on sagittal cryosections may not be sensitive to subtle differences in ECM composition or cell contractility that could be better detected with other approaches.

Additionally, dexamethasone may preferentially alter the juxtacanalicular tissue (JCT) region of the TM, as corticosteroid treatment has been reported to affect basement membrane material adjacent to the SC inner wall^46^. Consistent with this observation, prior studies from our group have demonstrated that the “outer” TM (proximal to the SC inner wall) is stiffer in DEX-treated mice compared to the “inner” TM^9^. The small size of the mouse eye, together with the spatial resolution of our force mapping approach with a spherical probe, limited our ability to resolve stiffness differences between TM sub-regions in the present study. Future investigations using larger animal models of corticosteroid-induced ocular hypertension or different approaches for measuring stiffness may permit layer-specific biomechanical measurements and thereby clarify how localized ECM and cellular mechanical alterations contribute to segmental outflow regulation.

Previous work measuring TM stiffness in HF vs. LF regions has focused on direct measurements of cells^20^ or ECM deposited by LF vs. HF-derived TM cells^24^. *In vitro* cell behaviors may not fully reflect the *in situ* state captured at the time of freezing, as TM cells may exhibit altered, potentially exaggerated, behaviors when removed from their native microenvironment. Further, finite element modeling based on *ex vivo* perfused human donor eyes has predicted greater stiffness in LF compared to HF regions, while corresponding AFM measurements did not reveal significant differences, despite the two methods being correlated overall in terms of stiffness estimation^47^. Taken together, these findings imply that regional mechanical differences may be highly context dependent.

Although overall TM stiffness did not differ between DEX-NP and CON-NP eyes, the lower minimum stiffness observed in LF regions of control eyes may reflect the presence of localized areas of softer tissue that facilitate aqueous humor outflow. Dexamethasone treatment may reduce the prevalence or extent of these softer regions, potentially contributing to increased outflow resistance and IOP elevation even in the absence of detectable changes in bulk tissue stiffness.

We also detected differences in labeling of ECM components following dexamethasone treatment. On average, fibronectin (FN) was elevated in DEX-treated eyes relative to controls, in agreement with prior studies^7,40,41,43,44,48,49^. Although FN levels were significantly elevated in the LF regions of DEX-treated eyes, correlation analyses with tracer intensity did not reveal consistent associations for either marker, indicating variability in protein levels across HF and LF regions. Importantly, FN is relatively compliant (effective modulus of order several kPa) compared to other ECM components, such as collagen, which can form matrices with stiffness of order MPa, so differences in FN deposition are not likely to drive significant changes in overall ECM stiffness. These findings suggest that our DEX-NP treatment increased outflow resistance in part through subtle changes in FN accumulation and cytoskeletal remodeling (α-SMA), which did not result in detectable changes in overall tissue stiffness with our AFM methodology.

In conclusion, this study demonstrates that dexamethasone-induced ocular hypertension in young mice subtly influences tracer distribution without significantly changing the proportion of high versus low flow regions. We found no detectable differences in trabecular meshwork average stiffness between HF and LF regions or between DEX-treated and control eyes using AFM force mapping, yet we did observe lower minimum stiffness values in the control LF regions relative to DEX-treated LF regions, as well as increased levels of FN in LF regions of DEX-treated TM. These molecular alterations are consistent with prior reports of glucocorticoid-induced remodeling and may contribute to elevated outflow resistance through mechanisms not captured by bulk stiffness measurements, such as localized changes in the juxtacanalicular region or dynamic cell-ECM interactions. Together, these findings underscore the importance of considering age and model-specific mechanisms in segmental flow studies and highlight the importance of integrating biomechanical and molecular assessments to better understand trabecular meshwork biology. Further research characterizing segmental flow differences across species, age-dependent effects, and detailed molecular differences in high and low flow regions will be critical for clarifying how segmental flow in DEX-induced ocular hypertension models translates to patients with ocular hypertension and POAG.

## Commercial Relationships Disclosure

C.A. Wong, None; A. T. Read, None; G. Li, None, A. Loveless, None; N.S. Fraticelli-Guzmán, None; A.J. Feola, None; T. Sulchek, None; W.D. Stamer, None; C.R. Ethier, None

