## Supplemental Files for "Segmental outflow and trabecular meshwork stiffness in an ocular hypertensive mouse model"

### Supplemental Materials

**Supplemental Table 1: Mouse eyes used in experiments and reasons for exclusion.** In cases noted as “off-target bead delivery”, fluorescent tracer beads perfused into the anterior chamber did not reach the TM but accumulated elsewhere (primarily the iris). “Needle displacement” indicates that the needle became dislodged during perfusion.

| Mouse | NP | Eye | Experiments (Reason for exclusion) |
|---|---|---|---|
| 1 | DEX-NP | OD | IOP + Tracer Perfusion + AFM |
| | | OS | IOP + Tracer Perfusion + AFM |
| 2 | CON-NP | OD | IOP + Tracer Perfusion + AFM |
| | | OS | IOP + Tracer Perfusion + AFM |
| 3 | CON-NP | OD | IOP only (Off-target bead delivery) |
| | | OS | IOP only (Off-target bead delivery) |
| 4 | DEX-NP | OD | IOP + Tracer Perfusion + AFM |
| | | OS | IOP + Tracer Perfusion + AFM + immunofluorescence |
| 5 | CON-NP | OD | IOP + Tracer Perfusion + AFM |
| | | OS | IOP + Tracer Perfusion + AFM + immunofluorescence |
| 6 | DEX-NP | OD | IOP + Tracer Perfusion + AFM |
| | | OS | IOP + Tracer Perfusion + AFM + immunofluorescence |
| 7 | DEX-NP | OD | IOP + Tracer Perfusion + AFM + immunofluorescence |
| | | OS | IOP only (Perfusion failure - needle displacement) |
| 8 | DEX-NP | OD | IOP + Tracer Perfusion + AFM + immunofluorescence |
| | | OS | IOP only (Perfusion failure- terminated early) |
| 9 | CON-NP | OD | IOP + Tracer Perfusion + AFM |
| | | OS | IOP only (Excluded for group size balance) |
| 10 | CON-NP | OD | IOP + Tracer Perfusion + AFM + immunofluorescence |
| | | OS | IOP only (Excluded for group size balance) |
| 11* | CON-NP | OD | IOP + Tracer Perfusion + immunofluorescence |
| | | OS | IOP only (Perfusion failure - needle displacement) |
| 12* | CON-NP | OD | IOP + Tracer Perfusion + immunofluorescence |
| | | OS | IOP + Tracer Perfusion |
| 13* | DEX-NP | OD | IOP only (arousal during perfusion) |
| | | OS | IOP only (arousal during perfusion) |
| 14* | DEX-NP | OD | IOP + Tracer Perfusion |
| | | OS | IOP + Tracer Perfusion |

*Eyes fixed by immersion prior to shipment

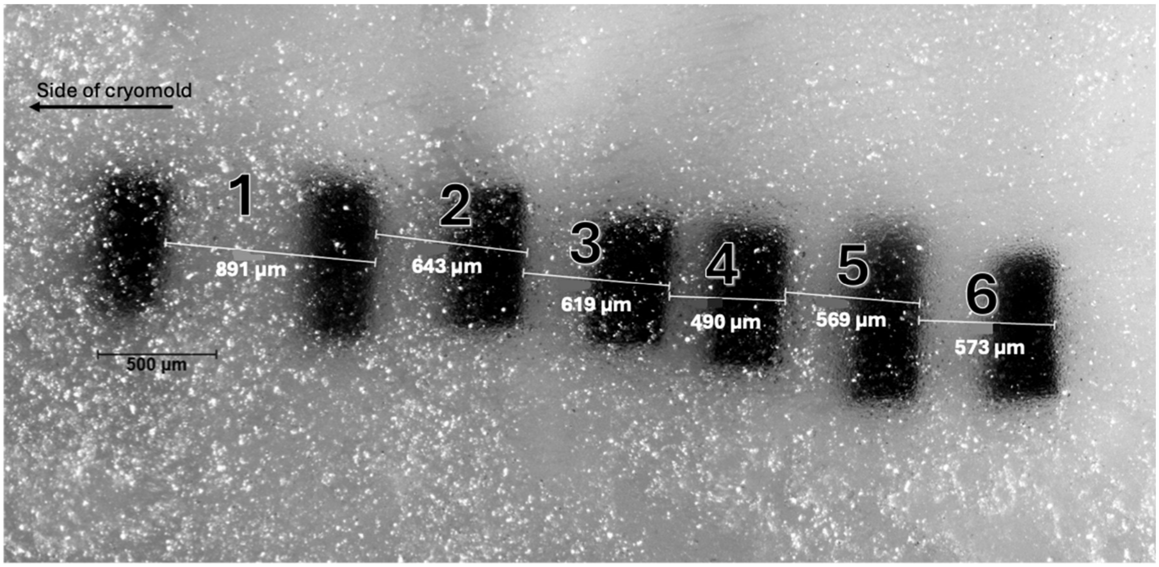


**Supplemental Figure 1**: **Cryomold with Phantom for Cryostat Calibration.** We first tested the accuracy of the cryostat used to produce sections of known, repeatable thickness so that cutting a predefined number of serial sagittal cryosections could accurately reach a pre-selected region within the tissue sample. For this purpose, we created a phantom, consisting of seven small rectangular pieces (~ 250 x 500 µm) of soft, thin plastic embedded in OCT (Supplemental Figure 1). This phantom was imaged on a Leica DMB 6 microscope with the 5x objective, and the distance between markers was measured using the microscope's software (LAS X). The phantom was then cryosectioned, keeping track of the number of sections and noting the value on the summation feature of the cryostat to estimate the distance cut away between markers. Numbered lengths correspond to values in

**Supplemental Table 2: Accuracy of Cryostat Sectioning Thickness.** Length numbers correspond to labeled distances in **Supplemental Figure 1.** Overall, the length estimated by the cryostat agreed well with distances measured microscopically, with an average error of approximately 5%.

| **Length #** | **Actual length between fiducial markers, as measured by the Leica DMB 6 microscope (µm)** | **Estimated length between fiducial markers, as measured by the Cryostar NX70 cryostat (µm)** | **Absolute Percent Error** |
|---|---|---|---|
| **1** | 891 | 950 | 6.6% |
| **2** | 643 | 630 | 2.0% |
| **3** | 619 | 650 | 5.0% |
| **4** | 490 | 430 | 12.2% |
| **5** | 569 | 580 | 1.9% |
| **6** | 573 | 600 | 4.7% |
| | | **Average error:** | **5.4%** |

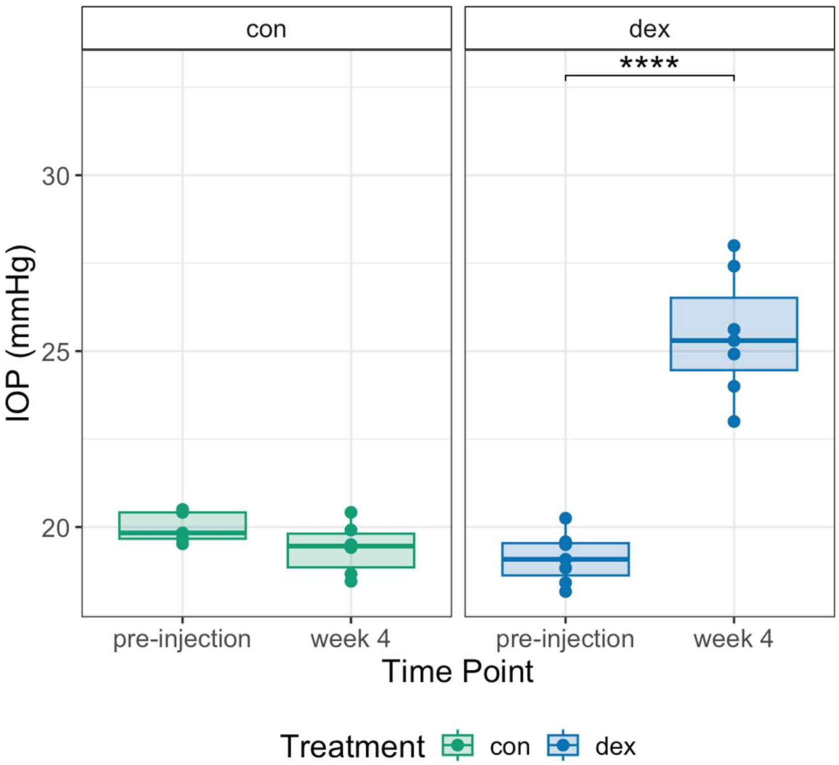


**Supplemental Figure 2: Change in IOP from baseline.** Intraocular pressure (IOP) at baseline and after 4 weeks of treatment, stratified by treatment group. Box plots show IOP measurements for each mouse at baseline and after 4 weeks of treatment. On average, DEX-treated eyes exhibited a significant increase of 6.35 ± 1.85 mmHg (mean: 33.3% increase) from baseline ($p < 0.0001$), while control eyes showed a modest, non-significant decrease of 0.45 ± 0.87 mmHg (mean: 2.19%).

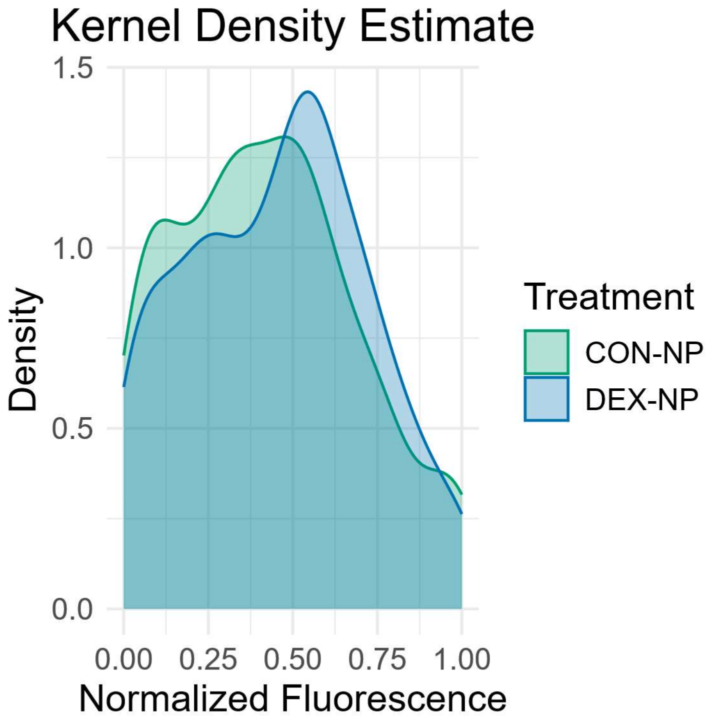


**Supplemental Figure 3: Kernel Density Estimate of Normalized Fluorescence Values.** Kernel density plot showing the distribution of normalized fluorescence intensity values from all DEX and CON-treated eyes, overlaid to highlight differences in distributions in the mid-range of values.

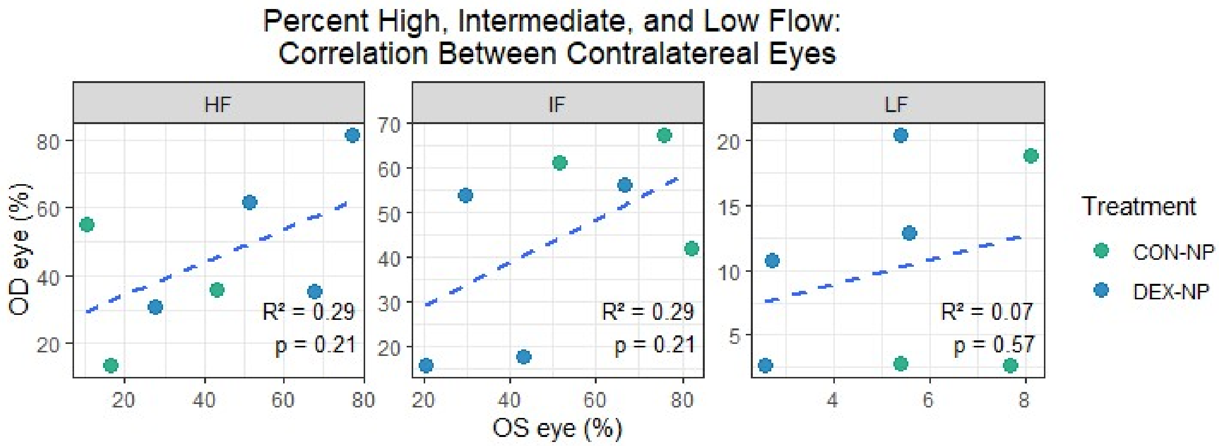


**Supplemental Figure 4: Correlation in Percentage of High, Intermediate, and Low Flow Between Contralateral Eyes.** For all mice where both eyes were successfully perfused (n=7), the percentage values for the OS eyes are shown on the horizontal axis, and the percentage values for the OD eyes are shown on the vertical axis. No significant correlation was observed for high, intermediate, or low flow percentages between contralateral eyes.

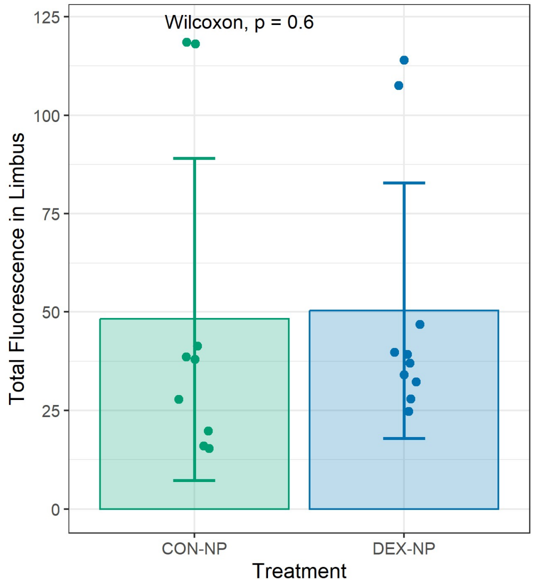


**Supplemental Figure 5: Total fluorescence in limbus of control and DEX-treated eyes.** The total fluorescence intensity in the limbus region (not normalized) did not show any differences between control and DEX-treated eyes. Each point represents a single eye, and bars represent the mean and standard deviation.
